# Ancient mitogenomes unravel massive genetic diversity loss during near extinction of Alpine ibex

**DOI:** 10.1101/2021.11.07.467348

**Authors:** Mathieu Robin, Giada Ferrari, Gülfirde Akgül, Johanna von Seth, Verena J. Schuenemann, Love Dalén, Christine Grossen

## Abstract

Population bottlenecks can have dramatic consequences for the health and long-term survival of a species. A recent bottleneck event can also largely obscure our understanding of standing variation prior to the contraction. Historic population sizes can be modeled based on extant genomics, however uncertainty increases with the severity of the bottleneck. Integrating ancient genomes provides a powerful complement to retrace the evolution of genetic diversity through population fluctuations. Here, we recover 15 high-quality mitogenomes of the once nearly extinct Alpine ibex spanning 8601 ± 33 BP to 1919 CE and combine these with 60 published modern genomes. Coalescent demography simulations based on modern genomes indicate population fluctuations matching major climatic change over the past millennia. Using ancient genomes, we show that mitochondrial haplotype diversity has been reduced to a fifth of the pre-bottleneck diversity with several highly differentiated mitochondrial lineages having co-existed historically. The main collapse of mitochondrial diversity coincided with human settlement expansions in the Middle Ages. The near extinction severely reduced the mitochondrial diversity. After recovery, one lineage was spread and nearly fixed across the Alps due to recolonization efforts. Contrary to expectations, we show that a second ancestral mitochondrial lineage has survived in an isolated population further south. Our study highlights that a combined approach integrating genomic data of ancient, historic and extant populations unravels major long-term population fluctuations.

## Introduction

The ongoing crisis of biodiversity loss is often referred to as the human-caused 6th mass extinction (Ceballos et al. 2015; Ceballos et al. 2020). But not only are species disappearing at an increasingly fast rate, anthropogenic pressures on population size and connectivity affect many more species, which are currently still at fairly good numbers. Hence, it becomes crucial to understand the scale and the genetic consequences of small population size and population fragmentation in the wild. Theory predicts progressive loss of genetic diversity, increased inbreeding and accumulation of deleterious mutations in small and isolated populations (Frankham et al. 2002; Hedrick and Garcia-Dorado 2016). Long standing theoretical and empirical work indicate the risk for reduced long-term survival and adaptive evolvability caused by low genetic diversity (Wright 1921; Keller and Waller 2002; Frankham 2010; Hedrick and Garcia-Dorado 2016; Hasselgren and Norén 2019). In this context, it is interesting that currently the protection status of a species seems to only poorly predict its level of genetic diversity (Díez-Del-Molino et al. 2018; Grossen et al. 2020). Furthermore, current genetic patterns may have different explanations. Low genetic diversity is usually explained by human induced population fragmentation and a sudden strong bottleneck, but also a long history of small population size and restricted gene flow may be its cause. Disentangling these two scenarios is important for proper species conservation measurements. Hence, this underlines the importance of taking into account the past demographic history of a species to predict its long-term viability. However, past bottlenecks (times of small population size) are not trivial to estimate. Recent methods based on coalescence theory such as PSMC (Pairwise Sequentially Markovian Coalescent, Li and Durbin 2011) and more recently MSMC (Multiple Sequentially Markovian Coalescent, Schiffels and Wang 2020) have been widely used to estimate the demographic history of species using contemporary DNA and investigate the impact of environmental changes on trajectories of past population size (Palkopoulou et al. 2015; Kozma et al. 2016; Kozma et al. 2018; Pečnerová et al. 2021). However, the study of recent DNA is expected to be limited in power due to uncertainty of past diversity, most of all after strong bottlenecks leading to significant loss of signal. Furthermore, estimates for the recent past, which was mostly affected by human presence on Earth, are not possible (Li and Durbin 2011; Nadachowska-Brzyska et al. 2016).

Ancient genomics is a promising state of the art method to solve this issue (Díez-Del-Molino et al. 2018). The study of ancient DNA (aDNA) allows quantifying population size over time and retracing changes of diversity through near extinction events thereby shedding light on the most important factors shaping current genetic diversity patterns. As expected, several aDNA studies focusing on pre-bottlenecked diversity suggested a major role for human impact on current patterns of genetic diversity (e.g. Casas-Marce et al. 2017). The Iberian lynx for instance, displayed a clear reduction in population size due to overhunting and divergence into two distinct subpopulations with drops to alarmingly low genetic diversity within a century (Casas-Marce et al. 2017). However, humans may not always be to blame. Studies on Musk ox and Kea interestingly identified environmental changes as the most important drivers of diversity loss suggesting prolonged histories of small population size (Campos et al. 2010; Dussex et al. 2015), while (Larsson et al. 2019) revealed a complex interplay of climatic and anthropogenic factors in arctic fox. There is also contradicting evidence and hence an ongoing debate on the role of humans in the Late Quaternary megafauna mass extinction (Koch and Barnosky 2006; Campos et al. 2010; Sandom et al. 2014; Lord et al. 2020; Stewart et al. 2021). Due to their imminent exposure to changing glaciation levels, demographic histories of species living in arctic or alpine habitats are expected to have been considerably affected by past climatic changes (Sommer 2020). While recent aDNA studies have mainly shed light on extinct arctic or alpine species (Palkopoulou et al. 2015; Gretzinger et al. 2019; Barlow et al. 2020; Lord et al. 2020; Ramos-Madrigal et al. 2021), less studies have investigated alpine species which survived the megafauna extinction event of the Pleistocene-Holocene boundary (e.g. Ureña et al. 2018).

An alpine species with currently low genetic diversity, high mutation load, high levels of inbreeding and signs of inbreeding depression, is the Alpine ibex (*Capra ibex*, Biebach and Keller 2009; Brambilla et al. 2018; Grossen et al. 2018; Bozzuto et al. 2019; Grossen et al. 2020). Historic records suggest that Alpine ibex were intensely hunted presumably since the 15th century and encountered their most severe bottleneck in the 19th century. The species survived near extinction in a single small population in a region which is today known as the Gran Paradiso National Park in Italy (Stüwe and Nievergelt 1991) before intense conservation efforts led to a fast recovery of the species during the 20th century. As predicted by theory, the severe bottleneck of close to 100 individuals left clear genetic footprints in contemporary populations (Biebach and Keller 2009; Grossen et al. 2018). Yet, the eradication of the species from almost its entire species range also eradicated signals of the past demography and it remains currently unclear what level of diversity was present before the near extinction. Due to the specific ecological needs of Alpine ibex (Grignolio et al. 2003; Grignolio et al. 2007), past populations may also have been small and isolated. And, given its current distribution in previously glacier-covered habitats, it is plausible that environmental changes in the late Pleistocene and Holocene also played a substantial role in shaping genetic patterns (Seersholm et al. 2020).

Here we combine diversity estimates covering the last eight millenia to get insight into the demographic and genetic history of the nearly extinct Alpine ibex. Taking advantage of published whole genome sequences, we quantify the effective population size of Alpine ibex over time and compare our observations with related species. The analysis of ancient and historic mitogenomes allows retracing changes of diversity and the demographic trajectory through the species near extinction. We quantify the current haplotype diversity in the light of past diversity and identify lost haplotypes and most affected gene regions. Finally, we investigate to what degree modelling based on current genetic data can recapitulate our new insights gained from pre-bottleneck sampling.

## Results

### Demographic history of Alpine ibex and related species

To explore past demographic fluctuations, we first took advantage of a published whole-genome dataset (Figure 1A) including Alpine ibex (N=29), Iberian ibex (N=4), Bezoar (N=6), Siberian ibex (N=2), Nubian ibex (N=2) and domestic goat (N=16) (Alberto et al. 2018, Grossen et al. 2020). Average depth of coverage ranged from 6.9 × to 46 × (Table S1). Past effective population size for each individual was estimated by applying the PSMC method (Li and Durbin 2011). The PSMC infers historical recombination events within a diploid genome and the time to the most recent common ancestor (TMRCA). The inferred population dynamics suggested comparable trajectories for all species under study with two rises in Ne estimates each followed by a Ne contraction (Figure 1B, Figure S1). The first contraction led to a local Ne minimum between 100 and 250 kya for all species except for the Iberian ibex (at ~75 kya). Ne estimates of Alpine and Iberian ibex followed a joint trajectory until approximately 200 kya (Figure 1B). This is earlier than previously estimated split times based on mitochondrial sequences (Acevedo and Cassinello 2009; 50 - 90 kya, Ureña et al. 2018). While the observed trajectories were similar among its relatives, Alpine ibex stuck out with a nearly flat trajectory of all individuals after the split from Iberian ibex, suggesting that their demographic history considerably differed from the other species with relatively low Ne estimates (Figure 1B). When compared to related wild species, mitochondrial nucleotide diversities suggested lower diversity in Alpine ibex, also for ancient specimens (Figure 1C).

**Fig. 1.**
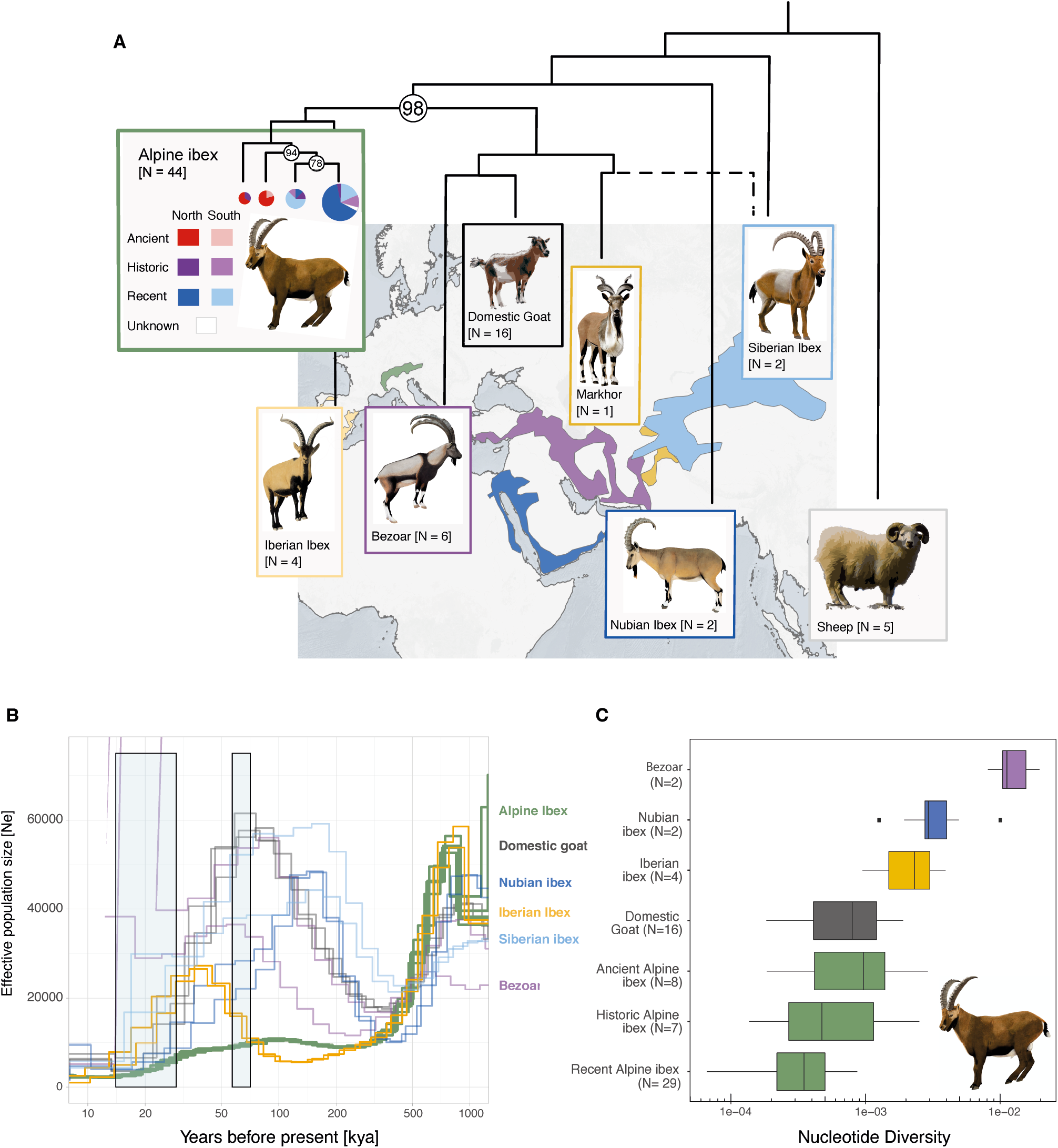
(*A*) *Capra* species distribution (except domestic goat) as stated by the IUCN and their phylogenetic relationship (Maximum likelihood tree, RAxML) based on 80 whole mitogenomes. Alpine ibex (*C. ibex*, N=44), Iberian ibex (*C. pyrenaica*, N=4), Bezoar (*C. aegagrus*, N=6), Siberian ibex (*C. sibirica*, N=2), Markhor (*C. falconeri*, N=1), Nubian ibex (*C. nubiana*, N=2) and the domestic goat (*C. hircus*, N=16) were included in the phylogeny. Sheep (*Ovis sp.*, N=5) were used as the outgroup. Nodes with Bootstrap support lower than 100 are explicitly stated and branches with a bootstrap support lower than 78 were collapsed. N indicates the number of mitogenomes used, dotted lines indicate a trans-species assignment. (*B*) Pairwise sequentially Markovian coalescent approach (PSMC) analysis based on Alpine ibex specimens from the Gran Paradiso National Park (N=6) and five related *Capra* species (N=2 per each species). The PSCM was constructed over 29 autosomes, a generation time of 8 years and a mutation rate of 3.568e-08 sites per generation was assumed. Marine oxygen isotope stage MSI 2 and MSI 4 are depicted in blue. (*C*) Mitogenome nucleotide diversity π per each *Capra* species with N>2 (Alpine ibex per sample age group) based on 15986 known sites and shown as boxplots. Siberian ibex are not included due to a cyto-nuclear discordance (see main text). The solid line indicates the median, the box spans from 25 % to 75 % of the interquartile ranges and upper and lower whisker spanning 1.5 * interquartile range.

PSMC analyses allow exploring estimates for past effective population size using recent genome-wide data (Li and Durbin 2011). However, changes of Ne estimates over time do not necessarily reflect actual species estimates of Ne, but can be confounded by population structure (overestimation of Ne) or restricted geographic distribution of direct ancestors (observed decrease of Ne estimates in the recent past, Olivier Mazet et al. 2015; Chen et al. 2019). Furthermore, from recent data alone, it is difficult to estimate historic diversity, most of all if a species went through a strong bottleneck and lost large parts of its past diversity and hence signal (O. Mazet et al. 2015). Therefore, to get a better understanding of the past demography of Alpine ibex, we here sequenced and analyzed 15 ancient and historic mitogenomes.

### Retracing the mitogenome diversity through a near extinction

To approximate the ancient genetic diversity of the Alpine ibex, we identified and collected samples which originated from up to several thousand years before and within the last strong bottleneck during the 19th century (Figure 2A). In particular, using shotgun sequencing, we analysed whole mitogenomes for seven ancient specimens (6601 ± 33 BCE to 1302 ± 26 BCE, six from caves, one from a glacier field) and eight historic specimens (1000 CE to 1919 CE) originating from museums and archeological excavations (Table S2, Figure 2A). Post-mortem-damage patterns were as expected for the respective sample age (Figure S2). The average endogenous DNA content was 62 % (ranging from 28.9 % to 92.6 %, Table S2). For a complete view on past and present diversity, we complemented our dataset with 65 additional published mitogenomes representing recent populations of six *Capra* species and domestic sheep (Figure 1A). The mean depth per individual ranged from 4.4 × to 2449 × for the historic and ancient samples and from 91 × to 7093 × for the recent samples (Table S2). Coverage along the mitogenome was >99.9 % in all samples and 2220 sites were found to be biallelic. 45 segregating sites were found among all Alpine ibex (including ancient, historic and recent specimens) after filtering for genotype quality and missingness.

**Fig 2.**
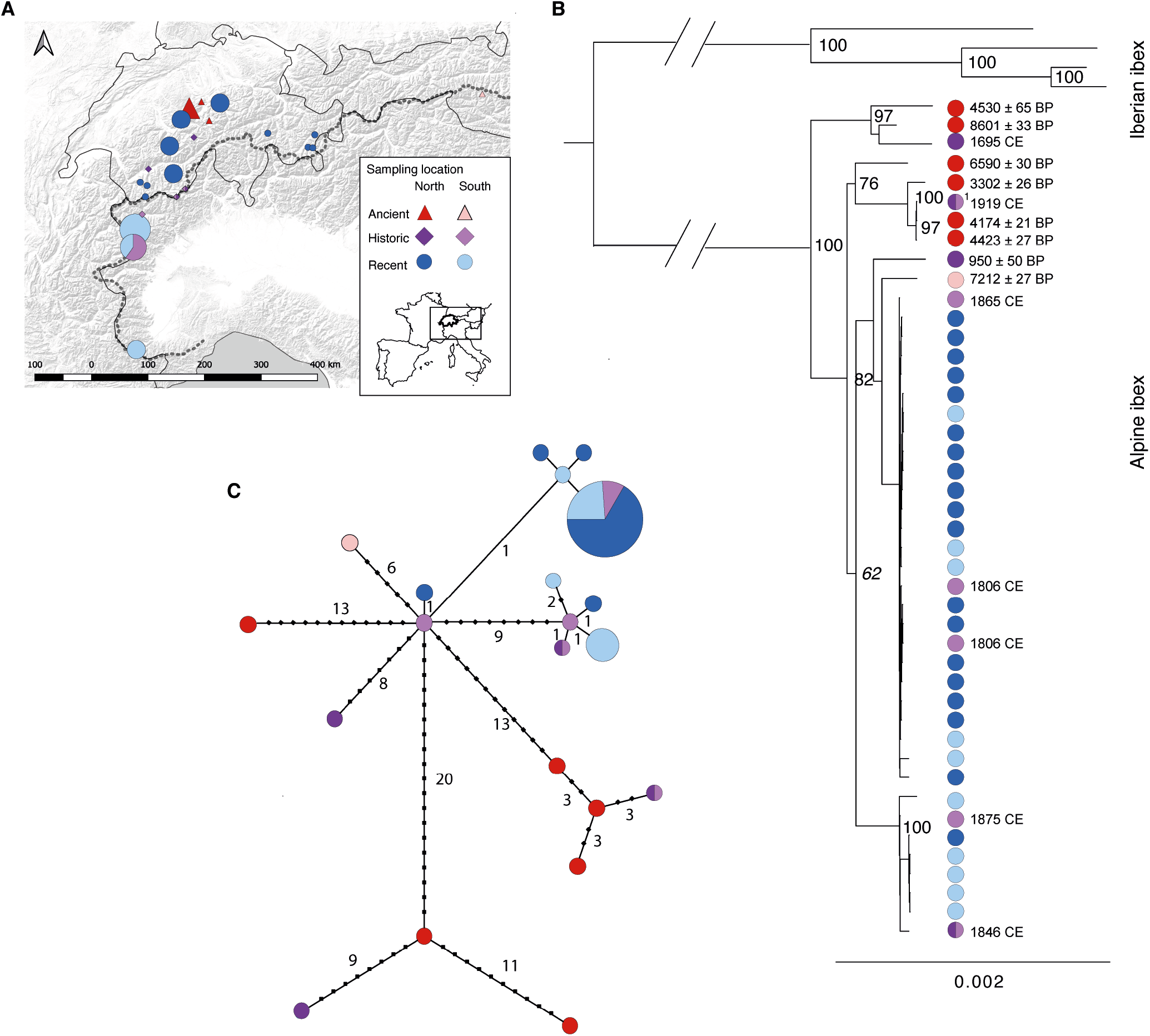
(*A*) Sample locations of ancient, historic and recent Alpine ibex specimens used for the study. Specimens which originated from the northern side of the main Alpine divide are colored in dark tones, whereas specimens originating from the southern part are colored in light tones. Specimens sampled on the Alpine ridge are shown in both color tones. The diameter of symbols indicates the sample size. Furthermore, triangles indicate ancient, diamond shapes historic and circles recent Alpine ibex specimens. (*B*) Maximum likelihood tree performed with a rapid bootstrap analysis and 100 repetitions for Alpine ibex and Iberian ibex. Major branches with a bootstrap value >60 are indicated. Color coding as in a. 1 indicates one individual with unknown origin. (*C*) Haplotype network including seven ancient, eight historic and 29 recent Alpine ibex samples. Little dots indicate mutational steps, size of pie chart indicates number of specimens representing the respective haplotype, color coding as described above (see Figure S4 for further detail).

We first investigated the phylogenetic relationship between recent, historic and ancient Alpine ibex specimens and other *Capra* species by performing a maximum likelihood phylogenetic analysis (Figure 1A). All Alpine ibex samples built a well-supported, monophyletic branch (bootstrap of 98), which was clearly distinct from the sister species Iberian ibex and all other *Capra* species in the tree (Figure 1A). The observed phylogenetic relationships among the other species (for instance bezoar and domestic goat were more closely related to Alpine ibex than Nubian ibex) confirmed previous findings based on mitochondrial DNA by Pidancier et al. (2006), except for one of the two Siberian ibex clustering with Markhor (Figure 1A, Figure S3). Assuming a constant substitution rate and a split time between Alpine and Iberian ibex of 57-92 kya, the most recent common ancestor of the sampled Alpine ibex was approximately 72 kya (Ureña et al. 2018). Among Alpine ibex, we found three distinct branches with moderate to high bootstrap support (Figures 1A, S3 and 2B). A further split separating the recent (and some of the historic) samples into two groups had only moderate support (bootstrap support of 78, Figure 1A). All recent Alpine ibex samples formed a monophyletic branch (bootstrap of 94) together with the historic southern samples (Gran Paradiso samples). This was expected, as all recent Alpine ibex populations derive from a single source population which survived the strong bottleneck in the 19th century in the Gran Paradiso National Park (Stüwe and Nievergelt 1991). The most basal branches of Alpine ibex were formed by ancient and historic northern samples (Figures 1A and 2B). The two historic specimens sampled very close to the main ridge of the Alps (Alpenhauptkamm, bicolour in Figure 2) either grouped with the northern ancient or southern historic and recent samples (Figure 2B).

Next, for a more comprehensive description of the intra-species diversity in Alpine ibex and to identify relationships in respect to their age and origin, we constructed a haplotype network of all Alpine ibex mitogenomes (Figures 2C, S4, Table S2). Allowing for one mutational step, we identified 14 distinct haplogroups with 21 haplotypes (Figure 2C). Most noticeable was the low haplotype diversity among the recent Alpine ibex populations with only two haplogroups (Figure 2C). The less frequent haplogroup was represented by six recent and two historic Alpine ibex individuals. Only one of these recent individuals originated from the northern part of the Alps, three from the southernmost population (Alpi Marittime) and two from the Gran Paradiso National Park (Figure 2C). One historic sample was from south of the Alps, one from the main ridge (bicolour, Figure 2C). The latter is assumed to be the last Swiss Alpine ibex individual shot in 1846 (personal communication Urs Zimmermann, game-keeper). The second major haplogroup was represented by a total of 26 individuals, both from south and north of the Alps and including three historic samples, all from the southern part of the Alps (Figure 2C). Within this haplogroup, 19 individuals were represented by a single haplotype (Figure 2C). Also two historic individuals from the Gran Paradiso National Park, source of all recent Alpine ibex populations, were assigned to the most abundant haplogroup. This may suggest that this haplotype was already common in this region during the time of the near extinction. All remaining historic and ancient specimens carried private haplotypes with up to 20 steps separating them from neighbouring haplotypes (Figure 2C), with no apparent geographic grouping. For instance, the two most divergent haplogroups were represented by ancient individuals found in close proximity (~20 km) and with similar time ranges (4423 ± 27 BP and 4530 ± 65 BP).

Compared to ancient and historic samples as well as all other *Capra* species, in recent Alpine ibex we found the lowest number of segregating sites, the lowest haplotype diversity and the lowest nucleotide diversity (all measured across the entire mitogenome, Table 1, Figure 1C). Furthermore, recent Alpine ibex were monomorphic in 8 out of 13 mitogenomic coding regions, which is in stark contrast to the ancient and historic specimens of the Alpine ibex and all other *Capra* species showing a maximum of 3 monomorphic coding regions (Table 1). A peak of diversity was found in the d-loop control region with the highest SNP density among the ancient samples (Figure S5). The recent samples only showed 9 segregating sites, of which 5 within coding regions. When comparing ancient and historic with recent diversity, a clear pattern of drift was observed, which led to the loss of most variants (Figure 3A). As expected by theory, variants more frequent among historic samples were also more likely to still be observed in recent samples. Variants observed at a frequency below 0.25 were generally lost while all variants with a historic frequency above 0.25 were also observed in recent populations (Figure 3B). Frequency among ancient samples did not necessarily predict frequency among recent samples (Figures 3A and S6). This is not unexpected, because the ancient samples were mostly sampled north of the Alps, while a large proportion of the historic samples originate from the source of all recent Alpine ibex populations (Gran Paradiso National Park).

**Table 1:**
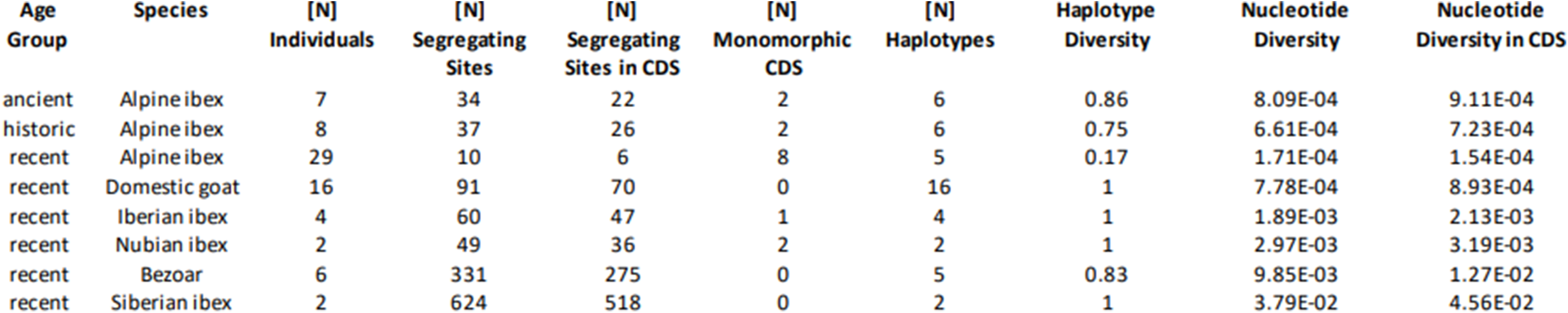
Summary statistics calculated based on 15986 mitochondrial sites per each *Capra* species with N>2 (Alpine ibex per sample age group). Siberian ibex are not included due to a cyto-nuclear discordance (see main text). Age group specifies the age of the samples: ancient (1302 ± 26 BCE to 6601 ± 33 BCE), historic (1000 CE to 1919 CE), and recent. Individuals [N] = Number of individuals, Segregating Sites [N] = number of segregating sites in the mitogenome, Segregating Sites CDS [N] = Number of segregating sites in coding regions, Monomorphic CDS [N] = number of monomorphic coding regions. Haplotypes [N] = Number of Haplotypes, Haplotype Diversity = Number of Haplotypes / Number of individuals, Nucleotide Diversity = Nucleotide diversity across whole mitogenome, Nucleotide Diversity in CDS = Nucleotide diversity in coding regions.

**Fig. 3.**
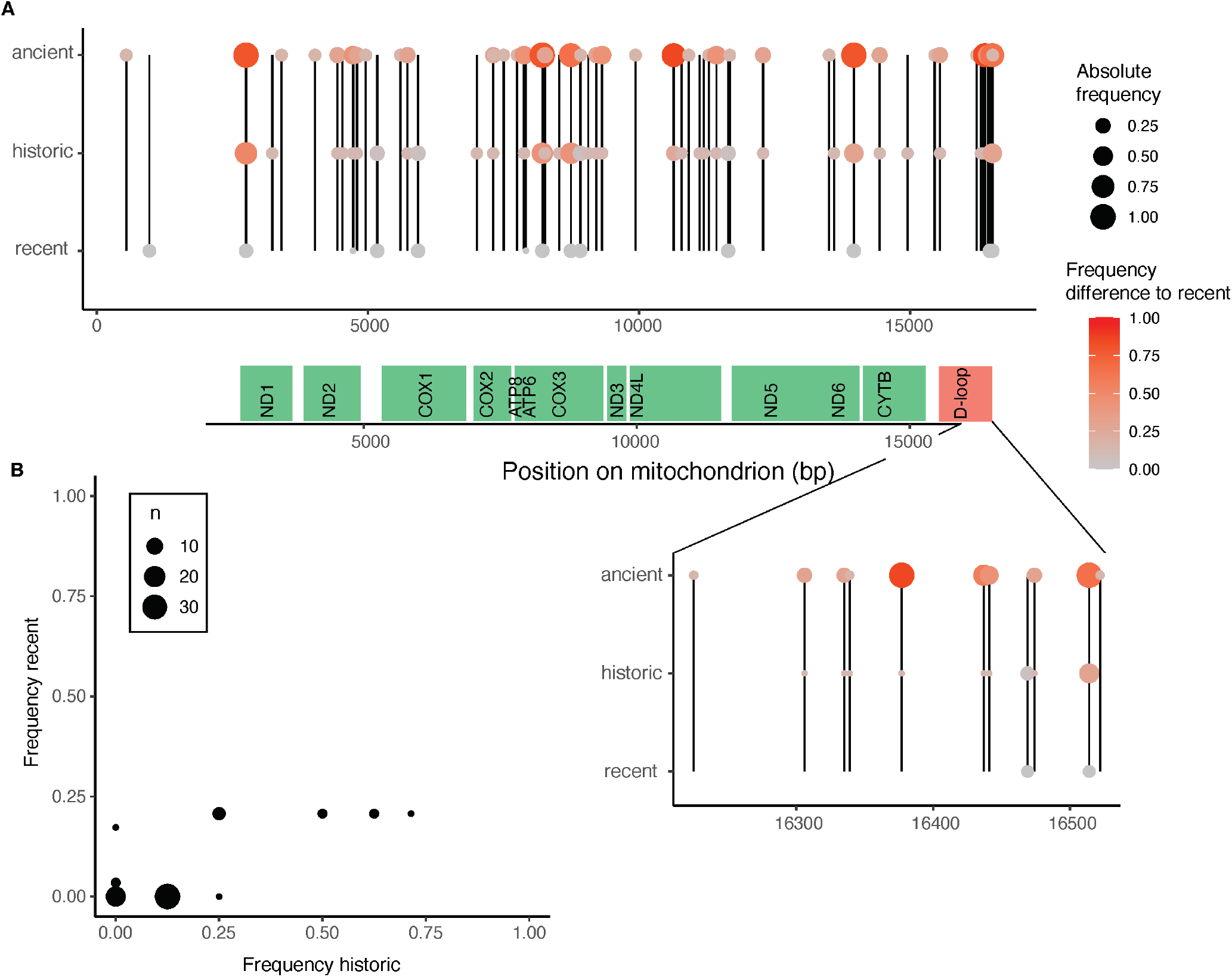
(*A*) Allele frequency of polymorphic sites along the mitogenome in ancient, historic and recent Alpine ibex. Coding regions are indicated in green. Vertical lines join the same site among sample age groups. Colors indicate frequency differences compared to recent Alpine ibex, with large differences indicated in red. Circle size represents absolute allele frequencies. The D-loop is zoomed in for better readability (*B*) Comparison of allele frequencies between recent and historic Alpine ibex. Circle size indicates the number of observations of a certain frequency combination.

We furthermore compared ancient, historic and recent Alpine ibex specimens by visualising segregating sites along the mitogenome as a circos plot, further illustrating the low genetic diversity among recent specimens (Figure S7).

### Stable demography before collapse during the last millennium

Historic records of Alpine ibex suggest a strong species bottleneck approximately 200 years ago. To determine the pre-bottlenecked demography of the species during the last interglacial cycle, we used the whole mitogenome data of all Alpine ibex individuals (seven ancient, eight historic and 29 recent mitogenomes) applying the Bayesian Skyline Plot approach (BSP) implemented in BEAST2 (Bouckaert et al. 2014). The estimates of past effective population size suggest nearly no fluctuations between 10 and 1 kya with an average Ne of approximately 4×10^3^ between 12 kya and 1 kya. These results suggest that until about 1 kya, Alpine ibex were at a relatively stable population size with no evidence for strong impacts from environmental changes or increasing anthropogenic influences. Evidence of significantly raising human activity along the European Alps can be found since 2.4 ky, intensifying during the last 1000 years (Boxleitner et al. 2017). The large-scale disappearance of the species is historically evident since the middle of the 16th century, which coincides with an elevated human population growth (Ziegler 1963; Stüwe and Nievergelt 1991; Head-König 2011). The estimated effective population size, inferred with the BSP, decreased nearly ten-fold to ~ 1.5×10^3^ during the last few centuries. This suggests that the Ne was relatively stable until the last millenium, when a rapid decrease in effective population size is evident.

## Discussion

In the present study, we retraced patterns of genetic diversity through a near extinction and compared our findings with current population samples. We analysed 44 high-quality mitogenomes spanning 8600 years and the current species range to get insight into the demographic and genetic history of Alpine ibex. We found a massive loss of haplotype diversity when comparing recent with pre-bottlenecked populations and identified 13 previously unknown haplotypes. Although higher a few thousand years ago, estimates of mitogenomic diversity of Alpine ibex were still lower than current estimates from related species. The analysis of published whole genome sequences revealed low long-term population size of Alpine ibex in comparison to related species. Hence, the genetic depletion of Alpine ibex was likely caused both by long-(environmental) and short-term (human-induced) factors. Our study shows how combining pre-bottlenecked with contemporary sampling provides a more comprehensive understanding of current patterns of diversity.

### Phylogenetic analysis confirms previously discovered cyto-nuclear discordance

Our phylogenetic analysis across species based on whole mitogenomes confirmed a cyto-nuclear discordance previously discovered based on cytochrome b sequences (Pidancier et al. 2006). Nuclear data places bezoar and domestic goat basal to Siberian and Nubian ibex and the latter two species closer to the two sister species Iberian and Alpine ibex (Pidancier et al. 2006; Grossen et al. 2020). However, our phylogeny based on mitogenomes placed bezoar and domestic goats next to Alpine and Iberian ibex (Figure 1C, S2). Wild goat species can interbreed and hence Pidancier et al. (2006) suggested mitochondrial introgression among ancestral taxa to explain this cyto-nuclear discordance. Interestingly, our data set includes a Siberian ibex, which, based on whole genome data, was clearly grouped with a second Siberian ibex (Grossen et al. 2020), but we here found that its mitochondrial haplotype clustered with the Markhor (Figure 1C). Hence, as a case example, it may have received mtDNA through introgression from Markhor. Cyto-nuclear discordance has been reported from a number of mammal species (Toews and Brelsford 2012).

### Alpine ibex show unique signals of demographic history compared to related species

The analysis of the recent Alpine ibex specimens using the PSMC method suggests a unique pattern of demographic history for Alpine ibex (Figures 1B, S1). The estimates of effective population size of the related ibex species show a relatively steep increase before further decrease during the last 100’000 to 200′000 years. However, estimates for Alpine ibex only barely increased again after the drop of Ne around ~250’000 years ago. This approximately coincides with a presumed, but strongly debated split time between the Alpine ibex and the Iberian ibex (Acevedo and Cassinello 2009; Ureña et al. 2018). Changes of coalescent-based Ne estimates over time can have different interpretations (Mather et al. 2020). An increase in Ne estimates can for instance demonstrate increased population structure (Heller et al. 2013; O. Mazet et al. 2015; Mather et al. 2020). A decrease in Ne estimates, as observed in most species for the more recent past, is often simply the result of recent ancestors having lived in closer and hence more closely related populations (Wakeley and Aliacar 2001). Hence, the rather flat Ne trajectory of Alpine ibex over the last 200′000 years may be explained by long-term local fidelity of the ancestor population. Alternatively, it could suggest relatively small species Ne over a long timescale. Similar declines of Ne estimates right after a lineage split were for instance found in Northern lions (de Manuel et al. 2020) and Amur leopards (Pečnerová et al. 2021) and were suggested to be the result of founding bottlenecks. The Late Pleistocene was determined by large ice shields covering large parts of Northern Europe and also the Alps (Seguinot et al. 2018). 115 to 13.7 kya determines the last large glaciation period. Considering the geographic distribution of Alpine ibex across the Alps (Figures 1A and 2A) and their specific ecological needs (Grignolio et al. 2003; rocks and alpine meadows, Grignolio et al. 2007), it’s plausible that the species was more strongly affected by these recent climatic changes than related ibex species. In accordance with prolonged times of relatively small population size is the observation of relatively low genetic diversity in ancient Alpine ibex when compared to the diversity found in related ibex species (Figure 1C). Past climatic changes have been suggested as driving forces for population size decline in a number of species including musk ox (Campos et al. 2010) and mammoths (Palkopoulou et al. 2015). And there is an ongoing debate on the role of climatic changes in the Late Quaternary megafauna mass extinction (Koch and Barnosky 2006; Sandom et al. 2014; Lord et al. 2020; Seersholm et al. 2020; Stewart et al. 2021).

### Massive loss of mitochondrial diversity

The analysis of more recent population size estimates using the skyline approach indicates a relatively stable effective population size until a drastic decline during the last few hundred years (Figure 4). As violations of the scenario of a single, isolated and panmictic population in coalescent-based demographic inferences can lead to spurious demographic signals, these results have to be interpreted cautiously (Heller et al. 2013). However, our findings are in accordance with historical records reporting a reduction of the Alpine ibex census size down to about 100 individuals at the beginning of the 19th century (Grodinsky and Stüwe 1987; Brambilla et al. 2020). The crash in population size coincides with known settlement expansions in the European Alps (Ziegler 1963; Chirichella et al. 2014). The main cause for the near extinction of Alpine ibex was most likely overhunting and increased competition with domestic ungulates (Stüwe and Nievergelt 1991; Acevedo and Cassinello 2009) as has been the case in a large number of species, in particular large mammals (Ripple et al. 2016). Similar erosion of mitochondrial diversity when comparing contemporary to pre-bottlenecked diversity was for instance found in the Iberian lynx (Casas-Marce et al. 2017).

**Fig. 4.**
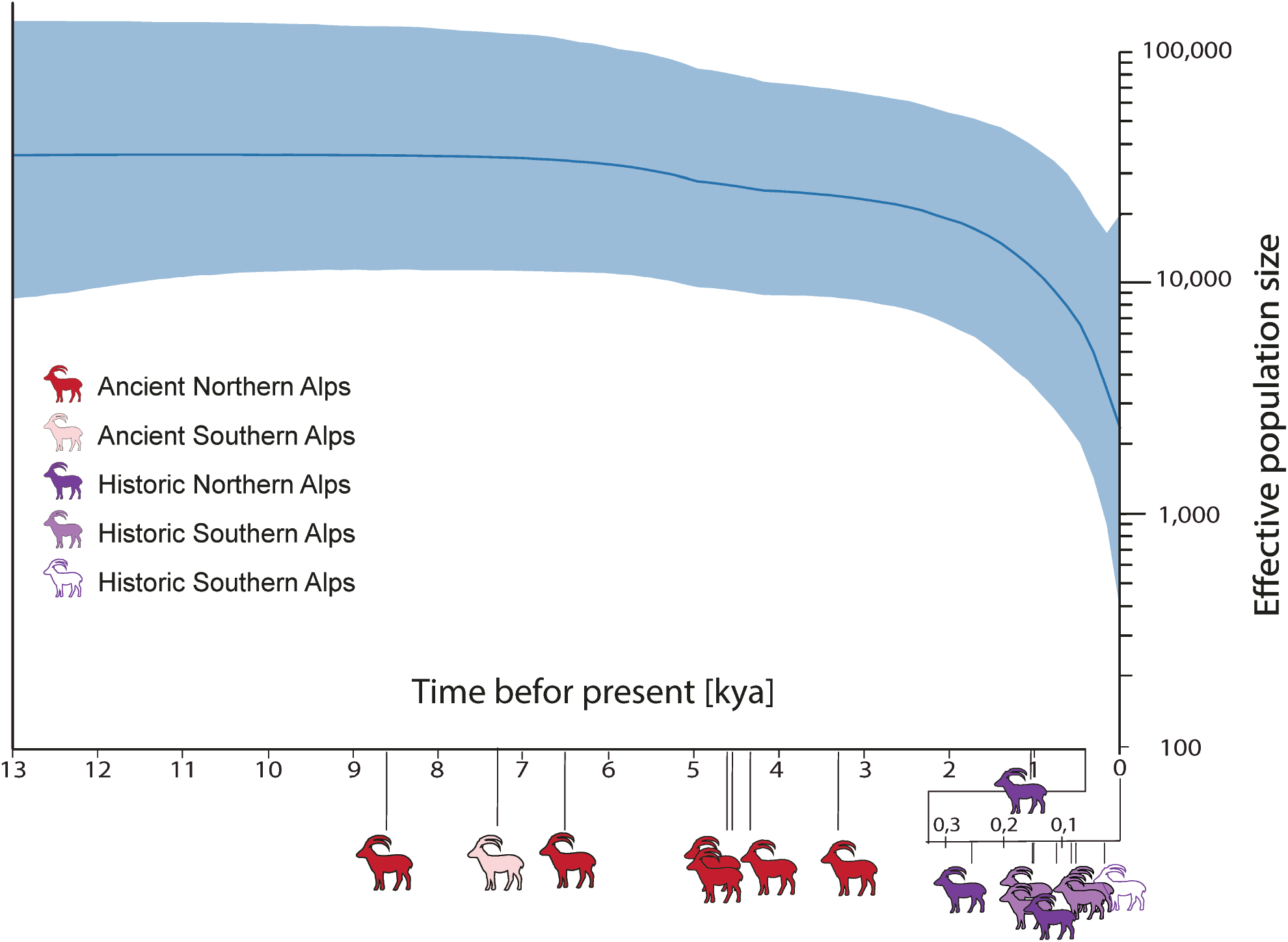
Bayesian Skyline-plot based on seven ancient (6601 ± 33 BCE) and eight historic (1000 CE to 1919 CE) as well as 29 recent Alpine ibex mitogenomes with a total of 16135 input sites. We used model averaging, incorporated a strict clock assumption, MCMC chain of 1.5*10^7^ samples and used a burn-in of 10%. Sample ages are indicated by ibex silhouettes along the time axis. Color tones indicate geographic origin (as in Figure 2).

The phylogenetic analysis revealed several divergent lineages among Alpine ibex. The deeper splits were (in terms of depth) comparable to what we observed among Iberian ibex. However, while no deeper splits were observed among recent Alpine ibex lineages, the Iberian ibex lineages were discovered (and hence still are existing) in recent populations. This is not surprising given the distinct demographic histories of the two species. Alpine ibex went through a very severe bottleneck (approximately 100 individuals) and are assumed to only have survived in one single population in Northern Italy (today the Gran Paradiso National Park, Stüwe and Nievergelt 1991). The Iberian ibex went through a less severe bottleneck (approximately 1000 individuals) and survived in several isolated populations. Although controversial, some of these populations were even classified as distinct subspecies by IUCN (Acevedo and Cassinello 2009; Groves and Grubb 2011: 224; Ureña et al. 2018; Sourp et al. 2020). As a consequence, the mitochondrial diversity remained larger in Iberian ibex than Alpine ibex confirming previous results based on genome-wide diversity measures (Grossen et al. 2018; Grossen et al. 2020).

Interestingly, some lineages with deeper splits were still represented by historic Alpine ibex suggesting that some of the ancient diversity was still present very close to the near extinction. This is also in accordance with nucleotide and haplotype diversity of historic samples being intermediate between ancient and recent Alpine ibex (Figure 1C, Table 1). Such findings give hope for other bottlenecked species with long generation times, because a prompt recovery from a bottleneck may save considerable diversity. But while the recent samples shared haplogroups with historic samples, none of the ancient haplogroups were observed among the recent samples. Accordingly, historic allele frequencies were correlated with recent allele frequencies, but ancient allele frequencies were only very marginally correlated with the recent ones. This is likely explained both by space and time. First, it is not surprising that the recent samples are genetically more similar to the historic samples just because there was less time in between for the mitogenomes to evolve. Second, a large proportion of the historic samples originates from the Gran Paradiso population, the source of all recent populations. More data will be needed to investigate how strong past population structure was, if the Alps formed a limit to gene flow and if Alpine ibex populations on both sides of the Alps were substantially differentiated from each other.

All the mitogenomic diversity observed among the recent samples was represented by only two haplogroups with nine mutational steps in between. Although 29 specimens is not a very large sample size, the sampling was explicitly chosen to represent the current species diversity (Biebach and Keller 2009; Grossen et al. 2018; Grossen et al. 2020; Kessler et al. 2020). Hence, it is unlikely that several common recent haplogroups remain undetected. As expected from the species history, individuals from Gran Paradiso (recent and historic) were found in either haplogroup. All recent individuals sampled North of the Alps (except for one) belonged to the most abundant haplogroup. The second haplogroup was represented by several recent and historic individuals, in particular from the population Alpi Marittime in Italy, a reintroduced Alpine ibex population occurring at the southern edge of the species distribution (Figures 1A and 2A). This population was reintroduced in the early 20th century, based on only about six founders. Accordingly, it shows high inbreeding and is highly divergent from all other existent Alpine ibex populations (Grossen et al. 2020; Kessler et al. 2020).

The most striking loss of diversity was observed in the d-loop (Figure 2). The d-loop is the regulatory region of the mitochondrial DNA and responsible for its replication and transcription (Nicholls and Minczuk 2014). It contains two hypervariable regions (HV-I, HV-II), which in humans have a 100 to 200 times higher mutation rate than the nucleus (Sharawat et al. 2010). Due to its high substitution rate, the d-loop can help to resolve differences between closely related individuals (Kundu and Ghosh 2015). The substantial differences in diversity in this region also underlines the severity of the most recent bottleneck which erased much of the rapidly evolving genetic diversity of the d-loop.

### Conclusions

We show a massive loss of mitogenome diversity and identify overhunting during the last centuries as the main cause of the low genetic diversity of contemporary Alpine ibex populations. However, the comparison with related species and the demographic modeling using whole genome data from recent populations suggests that Alpine ibex population size was reduced over prolonged times. Hence, although to a lesser extent, Alpine ibex demography was likely also affected by long term environmental processes such as glaciation. Our study underlines the value of a combined approach of ancient and historic mitogenomes, demographic modelling based on contemporary and related species data to understand past population fluctuations and their consequences on contemporary patterns of genetic diversity.

## Methods

### PSMC

To reconstruct the demographic trajectories of the Alpine ibex during the late Pleistocene, we incorporated a Pairwise Sequentially Markovian Coalescent (PSMC) approach (Li and Durbin 2011; Nadachowska-Brzyska et al. 2016). The PSMC infers historical recombination events within a diploid genome and facilitates differences in heterozygosity within one individual. It has improved statistical strength to infer deep-in-past events of coalescence by inferring the most recent common ancestor (TMRCA) and thereby the ancestral effective population size, depending on generation time, over the last 2*10^3^ to 3*10^6^ years (Li and Durbin 2011; Nadachowska-Brzyska et al. 2016; Mather et al. 2020). We used whole genome data, which has previously been published (Alberto et al. 2018, Grossen et al. 2020), representing Domestic goats (*Capra hircus*, N=3), Bezoar (*Capra aegagrus*, N=2), Nubian ibex (*Capra nubiana*, N=2), Sibiran ibex (*Capra sibirica*, N=2), Iberian ibex (*Capra pyrenaica*, N=2), and six Alpine ibex from the source population in Gran Paradiso, Italy. The whole genomic data of the domestic goat, Bezoar and domestic sheep are available through the NextSeq Consortium (see also Alberto et al. 2018). The data of the ibex species were obtained by Grossen et al. (2020). The reads were trimmed using Trimmomatic v.0.36 (Bolger et al. 2014) and subsequently mapped with bwa-mem v0.7.17 (Li 2013) to the domestic goat reference genome (ARS1, Bickhart et al. 2017). Duplicated reads were marked with MarkDuplicates from Picard v1.1301. The mean genome wide coverage was > 99.38 % for all samples and the average depth ranged from 6.9 × to 46 × (Table S1). To produce the input data set for the PSMC analysis, we followed the general pipeline suggested in Palkopoulou (2015). In detail, we used samtools mpileup (Li et al. 2009) in combination with the bcftools call command, keeping reads with a minimum mapping quality (-q) and minimum base quality (-Q) of 30 to produce an alignment. Next, we called consensus sequences using bcftools -c and performed a final filtering step using ‘vcf2fq’ from vcfutils.pl with options -d 5 and -D 34 (minimum and maximum coverage) and -Q 30 (minimal mean squared mapping quality). This pipeline has the advantage that the aligner does not assume Hardy-Weinberg equilibrium and does not rely on population frequencies for variant calling (Nadachowska-Brzyska et al. 2016). We then used fq2psmcfa to produce input data for psmc v. 0.6.5-r67, which was run with standard parameters as suggested by Li (2016). Specifically, the limit of TMRCA and the maximum number of iterations were left at the default values -t 15 and -N 25 respectively. Ne was inferred across 64 free atomic time intervals using the -p option with -p “4+25*2+4+6” which set the initial population-size parameter to four atomic time intervals followed by 25 parameters spanning two intervals followed by two parameters spanning four and six intervals respectively and allowed for 28 (1+25+1+1) free interval parameters. The psmcfa output was visualized in R, using a modified version of the plotPsmc.r script supplied by (Liu and Hansen 2017) with mutation rates of 2.23E-09 sites/year inferred for the siberian ibex (Chen et al. 2019). To explore the parameter space, we visualized the results with half and double the mutation rate (Figure S1A and S1C).

### Sampling

To analyse the ancient genetic diversity of Alpine ibex, we identified and collected samples which originated from before and during the last strong species bottleneck during the 19th century. We obtained a total of 22 (sampled in 2017) and four (sampled in 2020) specimens from Swiss museums, archeological institutions and cave excavations (Table S2). After screening for endogenous DNA content, a total of 15 samples were chosen for subsequent analysis. Samples originating from the last millenium are from now on referred to as historic samples. Their age was inferred by consulting registry entries of the museums of origin (or by C14-dating, Table S2) and ranged from 1000 CE (Common era) to 1919 CE. These specimens originated from Italy, France and Switzerland. The cave samples were AMS-C14 dated (between 8601 ± 33 BP and 3302 ± 26 BP) and are referred to as ancient samples. One specimen (Sample ID: Gro1) was found in a glacier field in the Austrian Alps and was AMS-C14 dated to 7212 ± 27 BP (Table S2).

To compare pre- and post-bottlenecked genetic diversity of the species, we included previously published whole-genome sequencing data representing recent populations of Alpine ibex (N=29), Iberian ibex (N=4), Nubian ibex (N=2), Markhor (N=1), Siberian ibex (N=2), Bezoar (N=6) and Domestic goat (N=16) (Alberto et al. 2018; Grossen et al. 2020). Additionally, previously published whole-genome sequencing data of five sheep individuals representing four species (Ovis sp.,Table S2, Alberto et al. 2018) was used as an outgroup for the phylogenetic analysis.

We produced a detailed map displaying the origin of recent, historic and ancient Alpine ibex specimens in Qgis v.3.0.2 (Figure 2A). For specimens where the exact location was not known (most of the historic samples), assigned coordinates are based on the region of origin (Table S2).

### Sample collection and DNA extraction

To ensure the accuracy of the aDNA results, established quality standards were incorporated (Rowe et al., 2011; 2005). To account for the usually minute quantity and degraded state of DNA in ancient samples, special care in sample handling was taken. (1) DNA extraction and library preparation were performed in a dedicated aDNA laboratory, (2) a one-way workflow from pre-PCR to post-PCR laboratory was integrated, (3) blank controls were used, (4) and used equipment was decontaminated with 7 % Sodium hypochlorite/NaClO and/or UV-irradiation.

For all sampling runs, DNA was extracted in a specialised clean-lab at the University of Zurich. In detail, we extracted DNA from teeth, skulls, long bones, petrous bones and horn material. During the sampling in 2017, surface sterilization of the samples was performed by washing the fragments in 1 % Sodium hypochlorite/NaClO for 1 min, followed by three washing steps with ddH2O. We extracted DNA from teeth, by detaching the cervix from the crown of the tooth with a precision drill (Dremel® 8200) at a low rotation rate of ca. 7000 rpm (speed level 10). To minimize friction, diamond cutting blades (Dremel® SC545) were used. Approx. 0.75 cm^3^ (horn) or 1 cm^3^ (bone or tooth material) were extracted with the Dremel. All 2017 samples were then ground to bone powder with an analytical mill (IKA™ A11 basic) and stored in micro test tubes at room temperature and under absence of light. For the 2020 sampling run, we sterilized the bone and horn surface with UV for 30 min per side, cleaned the surface with 7% Sodium hypochlorite/NaClO and removed the most outer part of the bone or horn with a dental drill and tungsten steel drill bits (Alpine Orthodontics, H1-014-HP). After removing an initial surface layer of bone, we extracted bone powder at low rotation rate.

To extract and purify mitochondrial and nuclear DNA from the bone and horn powder, we used a QIAquick® PCR Purification Kit and applied an established protocol for DNA extraction as described in (Dabney et al. 2013). The following modifications were made: we lysed 100 mg bone powder by adding 50 μl Proteinase K, 800 units/ml (New England Biolabs Inc.) to 950 μl EDTA and incubated at 37 °C overnight. The overnight digestion was centrifuged at 14 krpm for 5 minutes in a tabletop centrifuge, mixed with 10 ml binding buffer (without tween) and 400ul 3M sodium acetate. We centrifuged the resulting suspension at 1500rpm for 15 minutes. We stored the eluted samples at 4 °C.

### Library preparation and sequencing

State of DNA preservation and endogenous DNA content were inferred based on an initial screening run and subsequent shotgun sequencing on pools of double-indexed libraries. Double-stranded DNA libraries were built for all samples under strict precautions to avoid contamination. All pre-amplification steps for constructing the libraries were performed in a dedicated aDNA clean laboratory and non-template controls were included. For the 2017 samples, initial libraries were constructed for screening purposes at the Institute of Evolutionary Medicine at the University of Zürich following Kircher, Sawyer and Meyer (2012) and sequenced on an Illumina HiSeq 2500 system (1 lane, 200 cycles, paired-end) at the Functional Genomic Center Zürich (FGCZ). After screening, another set of libraries were built as individually double-indexed libraries optimised for aDNA (Meyer and Kircher 2010; Kircher et al. 2012), but here notably we corrected for post-mortem-damage by adding the uracil-specific excision reagent (USER™, New England Biolabs Inc.) during the blunt end repair step (for further detail see Supplementary Methods). The libraries were constructed in facilities of the Swedish Museum of Natural History in Stockholm, Sweden and deep sequenced on an Illumina HiSeq × system (one lane per individual, 300 cycles, paired-end) in a facility of the National Genomics Infrastructure (NGI), Sweden. Double-stranded DNA libraries for the 2020 samples (N=4) were built in a specialised aDNA clean laboratory at the Institute for Evolutionary Medicine, Zürich (Supplementary Methods) following the protocol described in (Meyer and Kircher 2010; Kircher et al. 2012). The sequencing was performed on one run of an Illumina NexSeq-500 in mid-output mode with 150 cycles (2*75+8+8) and sequenced at the Functional Genomic Center Zürich (FGCZ).

### Raw data analysis

Trimming of adapter sequences and read quality filtering was performed using Trimmomatic, ver. 0.36 (Bolger et al. 2014) with the following settings: ILLUMINACLIP:2:30:10:1:TRUE, LEADING:3, SLIDINGWINDOW:4:15, MINLEN:25. Trimmed reads were mapped using the Burrows-Wheeler Aligner (Li and Durbin 2009) BWA-MEM, ver. 0.7.15-r1142 to the *Capra ibex* mitochondrial reference genome NC_020623.1 (NCBI). Reads were sorted using samtools ver. 1.10 (Li et al. 2009). Duplicates were identified using MarkDuplicates from Picard (http://broadinstitute.github.io/picard, ver. 2.8.3). Summary statistics were produced using samtools v. 1.10. Post-mortem-damage of the 2017 samples was already confirmed at the screening step (Supplementary Figure S2) and the samples were USER-enzyme treated for the deep sequencing. Hence, no further post-mortem assessment was carried out for these libraries. Post-mortem-damage of the 2020 samples (Supplementary Figure S2) was assessed with mapDamage, ver. 2.0 (Jónsson et al. 2013). Base quality scores were corrected for post-mortem-damage by running mapDamage --rescale-only for the paired and unpaired reads separately. The published data representing all recent samples was also quality trimmed in Trimmomatic ver. 0.36 (Bolger et al., 2014) with settings ILLUMINACLIP:2:30:10 LEADING:5 TRAILING:5 SLIDINGWINDOW:4:15 MINLEN:50 and bam files were generated as described above.

We used HaplotypeCaller from GATK ver. 4.1.7 (Genome Analysis toolkit, Cooper and Poinar, 2000) to discover variant sites and produce g.vcf files for all recent, historic and ancient samples, before combining them and perform joint genotyping using GenomicsDBimport and GenotypeGVCFs. GATK Variantfiltration was used for hard filtering applying the following criteria to retain a site: Quality by Depth (QD) > 2.0, Mapping Quality (MQ) > 20.0, Mapping Quality Rank Sum (MQRankSum) >−3 or < 3,, Fisher Strand (FS) < 40.0, Strands Odds Ratio (SOR) < 5.0, ReadPosRankSum > −3.0 and < 3.0. We furthermore removed low quality genotypes with a genotype quality (GQ) below 20 (vcftools, Danecek et al. 2011).

We produced two mitogenome datasets, which differed in the number of individuals/species (Capra_all and Capra_ibex). For both datasets, we required a minimal genotyping rate per site of 90% (vcftools, Danecek et al. 2011). Hence, also the number of sites included differed between the two. The dataset Capra_all contained 80 individuals representing Alpine ibex (*N=44: 29 recent, 8 historic, 7 ancient*), Iberian ibex (*N=4*), Bezoar (*N=6*), Siberian ibex (*N=2*), Markhor (*C. falconeri, N=1*), Nubian ibex (*N=2*), domestic goat (*N=16*) and domestic sheep (*Ovis sp*., N=5). This dataset contained a total of 15986 of a possible 16157 sites. We furthermore produced a second dataset Capra_ibex, which only contained Alpine ibex specimens (*N=44: 29 recent, 8 historic, 7 ancient*). It was composed of 16135 out of a total of 16157 sites.

### Phylogeny

To explore the phylogenetic relationship among pre- and post-bottlenecked Alpine ibex and related species, we built two maximum likelihood trees. The trees were also used to confirm the species of our specimens, because the unambiguous morphological identification of small remains of Alpine ibex in relation to other *Capra* species can be challenging. Both trees were built using the dataset Capra_all, except that for the second tree, only Alpine ibex and Iberian ibex were included in order to allow a more detailed analysis of the diversity found among Alpine ibex. We used vcf2phylip to convert VCF to PHYLIP and defined all sheep (tree 1) or individual py.M518_sn (tree 2) as outgroup. We used the program RAxML, ver. 8.2.10 with the –GTRGAMMA model, which is a GTR model of nucleotide substitution under the ┌-model of rate heterogeneity (Prost and Anderson 2011; Stamatakis 2014). We performed a rapid bootstrap analysis with 100 repetitions and a search for the best-scoring Maximum Likelihood tree. The results were visualized with FigTree, v1.4.3 (Rambaut 2012) and edited in Adobe Indesign to improve readability.

### Haplotype networks

To infer and visualise the haplotype diversity among recent and past Alpine ibex, we built a haplotype network including all Alpine ibex specimens (ancient, historic and recent, dataset Capra_ibex). The tool vcf-to-tab (Chen 2014) and the Perl script vcf_tab_to_fasta_alignment.pl (Chen 2014) were used to convert the VCF format to FASTA. The FASTA sequences were then aligned using clustal-omega v. 1.2.4 (Sievers et al. 2011) which incorporates a clustalW algorithm. The resulting FASTA alignment was converted into PHYLIP using the Perl script Fasta2Phylip.pl (Hughes 1.2007). The R package TempNet (Prost and Anderson 2011) was then used to build the haplotype network using the incorporated TCS algorithm. TCS calculates an absolute pairwise distance matrix of all haplotypes and connects the haplotypes according to the parsimony criterion to minimize mutation steps between haplotypes. The resulting haplotype network was edited for improved readability in Adobe inDesign.

### Description of Mitogenome

We calculated mitochondrial diversity statistics with the R-package PopGenome v 2.7.5 applied to dataset Capra_all (Pfeifer et al. 2014). In PopGenome, by default, only sites genotyped in all individuals are retained. The gff3 file corresponding to the Alpine ibex mitogenome sequence NC_020623.1 was adapted for compatibility with the package {PopGenome} and was used with the command *set.synonym* to infer neutrality statistics. Neutrality statistics such as Tajmias’D were calculated with neutrality.stats (GENOME.class,detail=TRUE). We calculated the nucleotide diversity per each species (in Alpine ibex per sample age group) with the function GENOME.class@Pi/ and the haplotype diversity within each population with. Finally, we calculated nucleotide diversity within coding regions with the split_data_into_GFF_features and command. Using the dataset Capra_ibex, we furthermore constructed circos plots for the historic, ancient and recent Alpine ibex specimens to visualise the genetic variation along the mitogenome of the species with the R package circlize v.0.4.12 and a custom script programed in R, published in (Gu et al. 2014). SNP density among Alpine ibex (dataset Capra_ibex) along the mitogenome was computed using the option -- SNPdensity in vcftools (window size of 500 bp) and visualized in R {ggplot2}.

### Skyline plot

We inferred the demographic trajectories for the female Ne of Alpine ibex during the Holocene with a Bayesian Skyline approach (Drummond et al. 2005). This method facilitates Markov chain Monte Carlo (MCMC) sampling to infer a posterior distribution of effective population size through time by sampling directly from gene sequences. We used the dataset Capra_ibex, including all Alpine ibex samples. We performed the analysis with BEAST2 v2.5.2 (Bouckaert et al. 2014) and used model averaging by applying the BEAST Model Test as site model (Bouckaert and Drummond 2017), enabled estimation of mutation rate under strict clock assumption and ran a MCMC chain of 1.5*10^7^ samples and a burn-in of 10%. Change in initial mutation rates yielded similar results (not shown). Tracer v1.7.1 was used for visualization.

## Acknowledgements

We would like to acknowledge the different institutions for granting us access to ancient and historical Alpine ibex samples. The following institutions supplied samples for the aDNA analysis: Archäologischer Dienst Graubünden, Naturhistorisches Museum Bern, Nationalpark Hohe Tauern, Naturhistorisches Museum Winterthur, Natur-Museum Luzern, Stiftung Naturerbe Karst und Höhlen, Naturhistorisches Museum Basel, Naturhistorisches Museum Freiburg, Naturmuseum Olten and the Zoologisches Museum Zürich. Special thanks goes to Martin Trüssel for his engagement in collecting and organizing the ancient samples. We also thank Urlich Schneppat for valuable inputs and training regarding the petrous bone extraction. We further thank Daniel Wegmann for insightful discussions regarding the analyses and Daniel Croll for helpful comments on a previous version of this manuscript. We acknowledge the Functional genomics center Zürich for sequencing and the National Genomics Infrastructure in Stockholm funded by the Science for Life Laboratory, the Knut and Alice Wallenberg Foundation and the Swedish Research Council, and SNIC/Uppsala Multidisciplinary Center for Advanced Computational Science for assistance with massively parallel sequencing and access to the UPPMAX computational infrastructure.

This work was supported by the University of Zürich through a University Research Priority Program “Evolution in Action” pilot project grant, the Swiss National Science Foundation (31003A_182343 to C.G.), the University of Zurich’s University Research Priority Program “Evolution in Action: From Genomes to Ecosystems” (to V.J.S and G.F) and by FORMAS (grant numbers 2018-01640 and 2015-676 to L.D. and J.v.S). This study makes use of data generated by the NextGen Consortium (grant number 244356) of the European Union’s Seventh Framework Program (FP7/2010-2014).

## Data availability

Mitochondrial alignments produced for this project were deposited at the NCBI Short Read Archive under the Accession nos. SAMN21895033-SAMN21895047 (BioProject PRJNA767255).

## Author contributions

G.F. and C.G. conceived the project. M.R. acquired all samples. M.R., G.F, J.v.S. and G.A. carried out the sample and library preparation. L.D. and V.J.S. supported the project by enabling access to laboratory, consumables and sequencing facilities and giving advice. C.G. and M.R conducted all bioinformatic analysis and wrote the manuscript with input from all co-authors.

## Notes

### Competing Interest Statement

The authors have declared no competing interest.

